# Synaptic inhibition in the lateral habenula shapes reward anticipation

**DOI:** 10.1101/2021.01.21.427572

**Authors:** Arnaud L. Lalive, Mauro Congiu, Joseph A. Clerke, Anna Tchenio, Yuan Ge, Manuel Mameli

## Abstract

The nervous system can associate neutral cues with rewards to promote appetitive adaptive behaviors. The lateral habenula (LHb) contributes to such behaviors as rewards and reward-predictive cues inhibit this structure and engage LHb-to-dopamine circuits. However, the mechanistic understanding of reward encoding within the LHb remains unknown. We report that, in mice, acquisition of anticipatory licking in a reward-conditioning task potentiates postsynaptic GABAergic transmission, leaving excitatory synapses unaffected. Conversely, LHb-targeted manipulations of postsynaptic GABAergic function via pharmacological blockade or impairment of GABA_A_ receptor trafficking decrease anticipatory licking. Hence, inhibitory signaling within LHb enables the expression of appetitive behaviors.

## Introduction

The ability to anticipate appetitive events is common to all individuals. This process requires appropriate function of basal ganglia circuits and more specifically of dopamine signaling (Fiorillo et al., 2003; Romo and Schultz, 1990). Dopamine neuron contribution to reward encoding partly depends on upstream control stemming from the epithalamic lateral habenula (LHb). LHb neurons in both primates and rodents respond with phasic inhibition to unpredicted rewards and reward-predicting cues (Matsumoto and Hikosaka, 2009; Wang et al., 2017). These inhibitory responses mirror excitatory activity observed in midbrain dopamine neurons. Accordingly, lesion of the LHb impairs reward prediction error signals within dopamine neurons (Bromberg-Martin et al., 2010; Tian and Uchida, 2015). However, the mechanisms underlying LHb inhibition during reward encoding, and whether such process contributes to reward anticipation remains unknown.

We found that blocking GABA_A_ receptors in the LHb perturbs anticipatory licking emerging after a reward-conditioning task in head restrained mice. The expression of anticipatory licking occurred concomitantly with the potentiation of synaptic inhibition onto LHb neurons, while leaving intact excitatory neurotransmission. Finally, LHb-targeted, selective reduction of GABAergic transmission through manipulation of GABA_A_ receptor trafficking to the membrane impaired anticipatory licking.

## Results

### Contribution of LHb GABAergic transmission to appetitive behaviors

We trained mice in a classical Pavlovian conditioning paradigm in which an auditory cue (conditioned stimulus, CS) predicted an appetitive outcome (unconditioned stimuli, US, drop of water with 10% sucrose) throughout six consecutive days (Fig. 1a). Each trial started with an auditory cue (CS, 2 sec) followed by a 1-sec delay and the reward (US). Reward volume (20μl) and probability remained constant throughout training. Across days, mice rapidly developed robust anticipatory behavior, measured as an increase in lick frequency during the CS and delay period compared to baseline (Fig. 1a; black trace). This supports that mice learned the reward-predictive value of the CS.

**Fig. 1.**
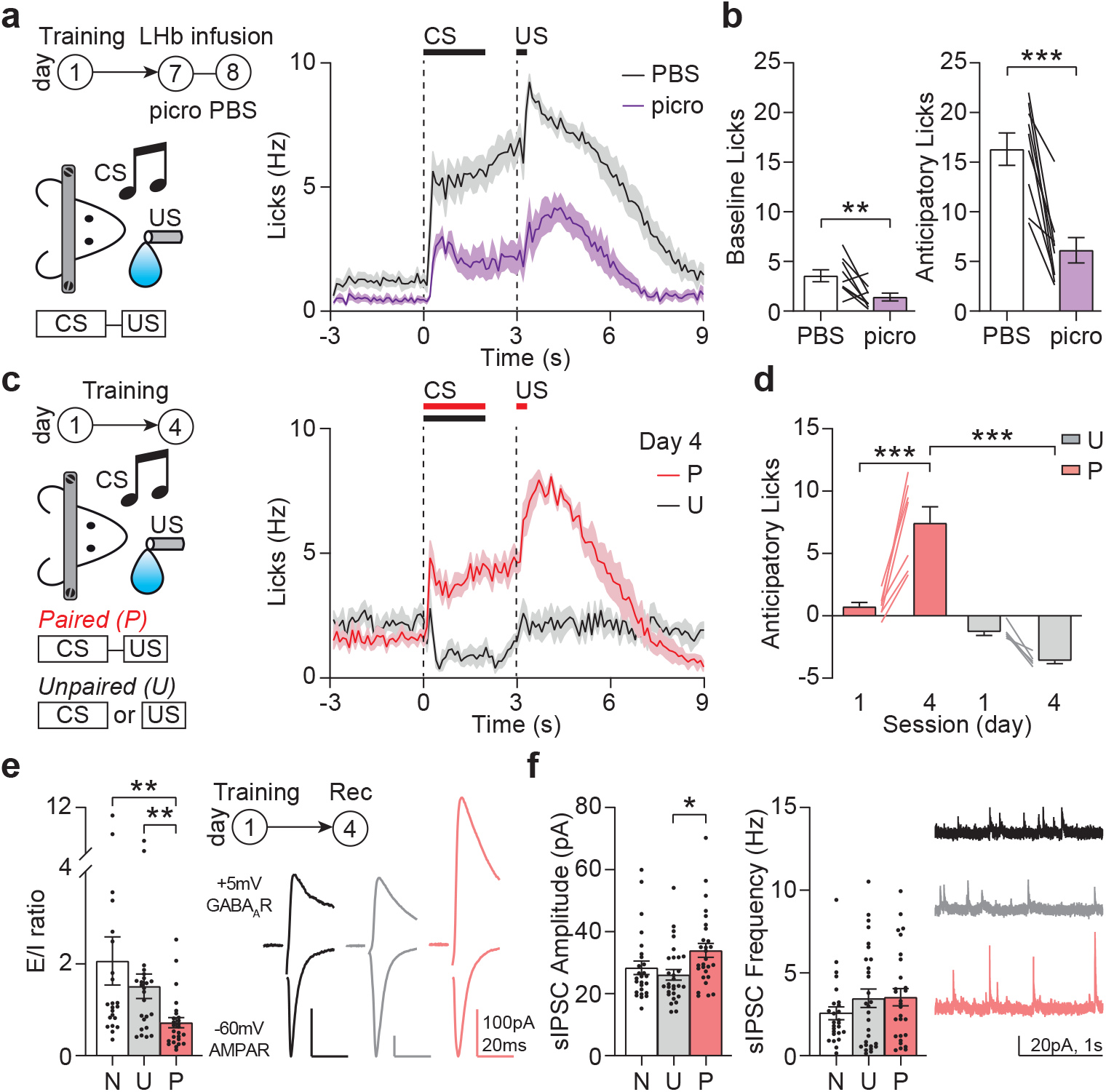
Reward anticipation potentiates LHb postsynaptic inhibitory transmission. a. Left: Pavlovian conditioning task design and timeline, combined with acute LHb infusion of picrotoxin or PBS. Right: Licking pattern of mice (n=9) infused with PBS (black) or picrotoxin (purple). b. Quantification of baseline licks (left, p=0.0056) and anticipatory licks (right, p=0.0003) after PBS or picrotoxin infusion. c. Left: Pavlovian conditioning task design. Right: Licking pattern after 4 days of training in paired (P, red, n=7) and unpaired (U, black, n=5) mice. d. Anticipatory lick quantification across training (2-way ANOVA, interaction F(1,10)=40.33, p<0.0001; Sidak’s P1 vs P4 p<0.0001; P4 vs U4 p<0.0001; U1 vs U4 p=0.1143). e. Excitation to inhibition ratio (E/I) and example traces from naive (N, n=22), unpaired (U, n=24) and paired (P, n=27) mice (Kruskal-Wallis p=0.0012; Dunn’s N vs P=0.0018; U vs P p=0.0154; N vs U p>0.9999). f. Left, spontaneous inhibitory current amplitude (N, n=26, U, n=29, P n=28; Kruskal-Wallis p=0.0113, Dunn’s U vs P n=0.0111; N vs P p=0.1134, N vs U p>0.9999). Right, spontaneous inhibitory current frequency (Kruskal-Wallis p=0.5861) and related example traces.

LHb neurons reduce their activity in responses to unpredicted and predicted rewards, supporting the implication of inhibitory signaling in this process (Matsumoto and Hikosaka, 2009). To examine the contribution of synaptic inhibition onto LHb neurons during stable anticipatory behaviors, we locally infused the GABA_A_ receptor blocker picrotoxin (100nl, 0.25mM) after an initial conditioning training (Fig. 1a). Blockade of GABA_A_ receptors acutely decreased anticipatory licking. Decreases were also observed in baseline and reward maximum lick rate. Importantly, total reward volume consumed remained identical (not shown). This effect promptly disappeared and high anticipatory licking was observed after control PBS infusion in the same mice 24 hours later (Fig. 1a, b). Retrobeads delivery at the end of the experiments enabled LHb targeting validation (Supplementary Fig. 1a). This indicates that GABAergic transmission shapes anticipatory licking when reward-learning processes are established.

### Concurrent potentiation of synaptic inhibition and anticipatory behavior

The anticipatory licking emerges from learning processes allowing the association between initially neutral cues and rewards (Cohen et al., 2012). Notably, long-term plasticity of synaptic transmission represents a substrate of various forms of learning, including cue-reward learning (Stuber et al., 2008; Tye et al., 2008). To test whether reward anticipation involves plasticity of inhibitory transmission in the LHb, we trained a group of mice in the Pavlovian conditioning paradigm (paired, P) and designed a separate control group where the CS and US were presented in a random order (unpaired, U, Fig. 1c). Only mice from the paired group showed significant increase in anticipatory licking (Fig. 1c, d). After the last training session (Day 4), we prepared acute slices containing the LHb. We probed excitatory and inhibitory transmission by holding LHb neurons at −60mV and +5mV, respectively. Currents were blocked by NBQX at negative potential, and by picrotoxin at positive potential, validating the isolation of selective AMPA receptor and GABA_A_ receptor postsynaptic currents (EPSC, and IPSC respectively; Supplementary Fig. 1b). Recordings revealed a reduction in EPSC to IPSC ratio (E/I) from paired mice compared with unpaired or naïve animals (Fig. 1e). To identify the origin of the decrease in E/I, we monitored spontaneous excitatory and inhibitory postsynaptic currents (sEPSC, sIPSC) from the same neurons. We found that sIPSCs amplitudes were significantly larger in the paired group, with no change in frequency or paired-pulse ratio (PPR) of evoked IPSC. In contrast, we found no difference in sEPSCs or EPSC-PPR across groups (Fig. 1f, and Supplementary Fig. 1c, d, e). Altogether, these findings identify postsynaptic strengthening of synaptic inhibition, but not excitation, onto LHb neurons, as reward anticipatory behavior is established.

### GABAergic synapses manipulation during anticipatory behavior

Manipulation of GABA signaling, and precisely of GABA_A_ receptor function has been proven challenging in light of the multitude of subunits of these proteins and their complex interactive protein network. Furthermore, knowledge about GABA_A_ receptor composition, dynamics and plasticity within the LHb is elusive (Meye et al., 2013). The Cleft lip and palate transmembrane protein 1 (Clptm1) is a GABA_A_ receptor-associated protein capable to interact with all GABA_A_ subunits and promote receptor trapping in the endoplasmic reticulum (Ge et al., 2018). Indeed, overexpression of Clptm1 reduces synaptic inhibition without altering excitatory neurotransmission in the hippocampus. We reasoned that viral-based Clptm1 overexpression in LHb (Fig. 2a) would be ideal to reduce postsynaptic GABA_A_ receptor expression and test whether this shapes anticipatory licking, thereby causally linking synaptic inhibition and behavior.

**Fig. 2.**
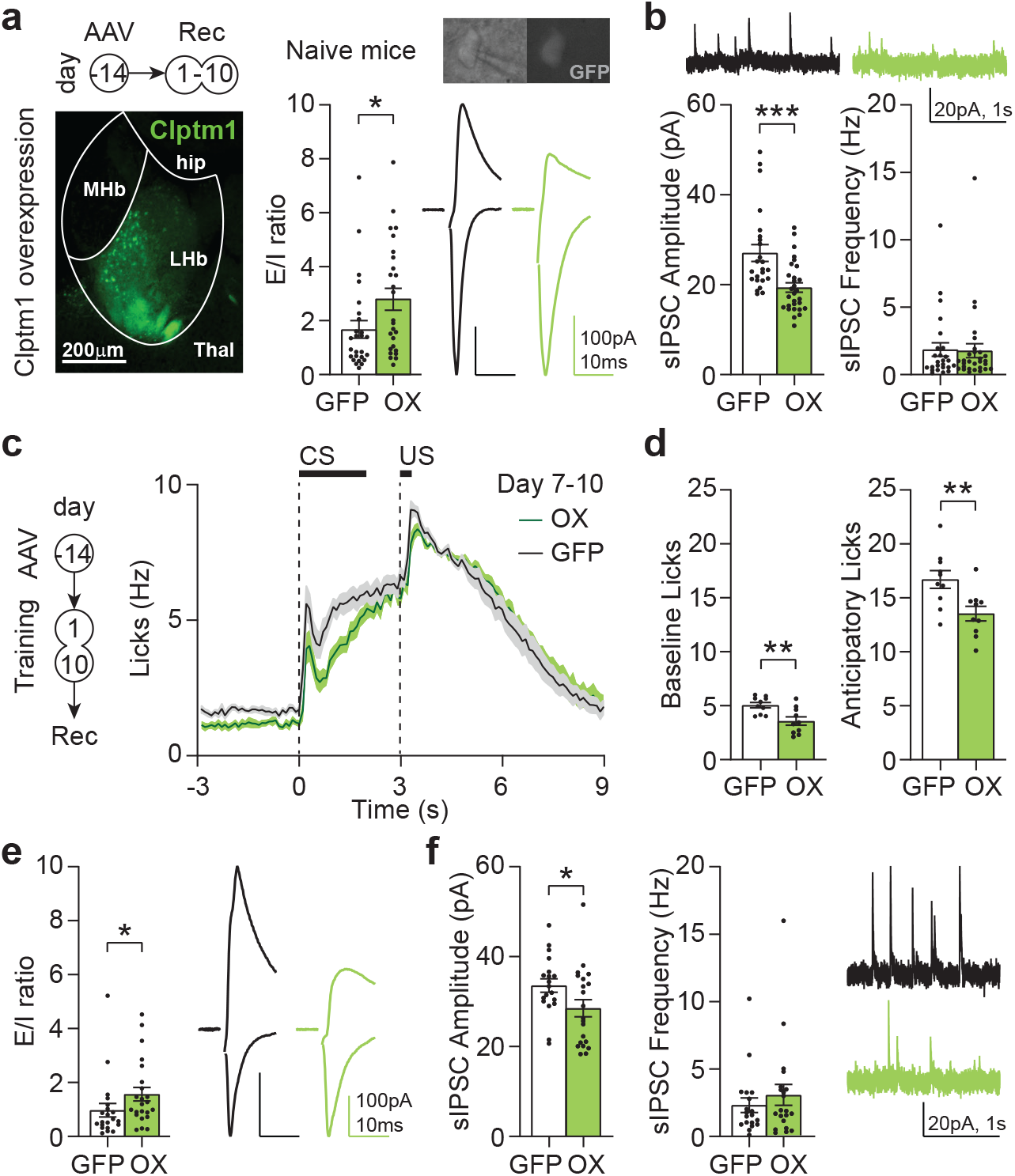
Postsynaptic GABA_A_ receptor downregulation in the LHb decreases reward anticipation. a. Left: Timeline and example picture of rAAV9-YFP-P2A-Clptm1 overexpression in the LHb. Right, E/I and example traces in control (GFP, n=26) versus Clptm1 overexpression (OX, n=26) in naive mice (p=0.0205). Inset, pictures showing targeted recording of GFP-positive neuron in LHb slice. b. Spontaneous inhibitory currents amplitude (left) and frequency (right), with example traces (GFP, n=24; OX, n=29; Amplitude p=0.0003; Frequency p=0.8465). c. Left: Timeline for Clptm1 overexpression, Pavlovian conditioning and recordings. Right: Licking pattern of control (GFP, n=10) or Clptm1 overexpression (OX, n=10) animals. d. Quantification of baseline licks (left, p=0.0089) and anticipatory licks (right, p=0.0115). e. E/I and example traces in neurons from control (GFP, n=21) or Clptm1 overexpression (OX, n=23) mice after Pavlovian conditioning (p=0.0157). f. Spontaneous inhibitory current amplitude (left) and frequency (right), with example traces, after Pavlovian conditioning (GFP, n=19; OX, n=21; Amplitude p=0.0472; Frequency p=0.5735).

Mice injected with rAAV9-YFP-P2A-Clptm1 to co-express YFP and Clptm1 in LHb presented larger E/I compared to control mice (infused with rAAV8-GFP control virus), along with a decrease in sIPSC amplitude but not frequency (Fig. 2a, b). Clptm1 overexpression did not alter the PPR of IPSCs nor excitatory neurotransmission (Supplementary Fig. 2a, b). Therefore, Clptm1 represents an appropriate tool to specifically decrease postsynaptic GABA_A_ receptors expression and weaken inhibitory transmission. Similarly to mice infused with picrotoxin (Fig. 1a, b), animals overexpressing Clptm1 exhibited reduced baseline and anticipatory licking when conditioning was established, compared to the control group (Fig. 2c, d). After the last behavioral session, the same mice were used for slice recordings. As predicted, E/I remained significantly larger in the Clptm1 overexpression group compared to controls (Fig. 2e). Similarly, sIPSC amplitude was smaller in the Clptm1 group while presynaptic parameters and excitatory synaptic transmission remained unaffected (Fig. 2f and Supplementary Fig. 2c, d). Altogether, this establishes causality between synaptic inhibition within the LHb and the maintenance of anticipatory licking in a reward conditioning task.

## Discussion

We found that the LHb carries information relevant for reward anticipatory behaviors. Namely, potentiation of synaptic inhibition onto LHb neurons contributes to anticipatory licking after reward conditioning.

A large body of information exists defining the LHb as an antireward center and mediator of aversive behaviors, processes mostly driven by LHb excitatory transmission (Lecca et al., 2017; Lecca et al., 2020; Proulx et al., 2014; Trusel et al., 2019). Our results highlight a new mechanism for a specific LHb function, namely the contribution of inhibitory transmission in shaping reward anticipation. LHb reduced activity in reward encoding has been suggested in rodents and monkeys, where rewards and reward-predicting cues drive LHb neuronal inhibition, presumably through GABA_A_ receptors (Matsumoto and Hikosaka, 2009; Wang et al., 2017). Here, we formally demonstrate the importance of postsynaptic GABA_A_ receptor transmission in shaping reward anticipatory behavior. Additionally, we provide a synaptic mechanism (postsynaptic GABA_A_ receptor potentiation) likely explaining the increase in calcium inhibition observed as mice learn a cue-reward association (Wang et al., 2017). The infusion of GABA_A_ receptors blocker and the Clptm1-mediated reduction in GABA_A_ receptor transmission suggest that synaptic inhibition scales with reward anticipatory behavior. Interestingly, both manipulations also affected baseline lick rate, outside CS or US presentation. It is possible that these licks still represent a form of reward anticipation, as the action of licking is directly linked to reward intake.

From a mechanistic point of view, the establishment of a synaptic potentiation concomitantly with reward anticipatory behavior may promote a stable tonic synaptic inhibition scaling down the activity of LHb. This may have repercussions on connected midbrain circuits. Accordingly, recordings in awake monkeys suggest that LHb activity during reward anticipation controls downstream tonic dopamine signaling (Bromberg-Martin et al., 2010). Whether inhibitory transmission also encodes rewards of different nature, or compute reward probability remain however open questions.

Photometric analysis of calcium fluorescence signals and single unit recordings in rodents suggest the existence of distinct LHb neuronal populations that respond differentially during reward anticipation (Wang et al., 2017). Some LHb neurons show phasic inhibition to CS and US, while others exhibit CS-driven inhibition followed by excitation. Our electrophysiological experiments reveal large variance across synaptic recordings indicative of non-homogeneous synaptic strength. Altogether, these data support the existence of specialized neuronal populations that may arise from connectivity or genetic differences (Hashikawa et al., 2020; Lecca et al., 2017; Wallace et al., 2020). Probing genetic, circuit, and functional properties may help to identify distinct LHb cell-types underlying aspects of reward encoding (Wall et al., 2019).

The manipulation of Clptm1 to control the efficacy of synaptic GABA_A_ receptor transmission in the LHb opens to several considerations. The finding showing Clptm1 overexpression-driven reduction in GABA_A_ function indicates that endogenous receptors are sensitive to Clptm1 control. The efficacy of Clptm1 in tuning inhibitory transmission in the hippocampus and in the LHb suggests that Clptm1 is an efficient tool to limit forward trafficking of the majority of GABA_A_ receptors throughout the brain (Ge et al., 2018). The relevance of GABA transmission in the LHb, for reward anticipation but also for depressive states in drug abuse and mood disorders makes Clptm1 an attractive target for intervention in the pathological context (Meye et al., 2016; Shabel et al., 2014). Understanding the temporal scale and precise mechanisms by which GABA_A_ receptor transmission potentiates concomitantly with anticipatory licking, and whether Clptm1 is part of this process remains an important aspect for future investigation.

In conclusion, we showed here that GABAergic synaptic transmission shapes reward anticipation, contributing to the knowledge of how the LHb instructs cardinal behaviors in physiological and pathological conditions.

## Methods

### Surgical procedures

All procedures were in accordance with the veterinary office of Vaud (Switzerland, license VD3172). Animals (C57BL6/6JR, Janvier, 7-12 week-old, males) were anesthetized with an i.p. injection of ketamine (150 mg per kg) and xylazine (100 mg per kg) and were placed on a stereotactic frame (Kopf). The scalp was opened and bilateral holes were drilled above the LHb. Bilateral injections of 200nl were performed through a glass needle at a rate of approximately 100 nl min^−1^ (coordinates from bregma in mm : AP −1.2, ML 0.45, DV from skull 2.6). The injection pipette was withdrawn from the brain 5 min after the infusion. When required, a stainless steels headbar was implanted on the skull. The skull was scraped clean and covered with a layer of Cyanoacrylate glue (Vetbond, 3M). The headbar was lowered to touch the skull over lambda, then secured to the skull with a layer of dental adhesive (C and B Metabond, Parkell) followed by dental cement (Jetkit, Lang).

### Viral constructs

For Clptm1 overexpression, we used an AAV9-hSyn-YFP-p2a-Clptm1* (gift from Prof. Ann Marie Craig) and an AAV8-hSyn-GFP as control (Zuerich University Vector Core).

### Awake infusions in LHb

For awake drug infusions, LHb sites were marked with sharpie pen on the skull and covered with a drop of silicone elastomer (Kwik-Cast; WPI). Mice were then implanted with a headbar (see above). Only a thin layer of cement was applied above the marked VM sites. One day before the experiment, mice were anesthetized, dental cement and silicone above LHb were removed, and holes were drilled on the marked LHb locations. The craniotomy was then covered in silicone elastomere and mice were given 24h to recover. For drug infusions, mice were headfixed, levelled flat, and a glass pipette mounted on a microinjector was lowered in the LHb to deliver 100 nl of picrotoxin (0.25mM in saline) or PBS (1%) over 2 min. The pipette was removed 2 min later and the same procedure was repeated in the other hemisphere. The craniotomy was covered in silicone elastomer and mice were immediately transferred to the behavior boxes for training. After the last behavior session, mice were infused with 20 nl of retrobeads in each side before cardiac perfusion with PFA.

### Pavlovian Conditioning

For Pavlovian reward conditioning, mice were headfixed in a 3-cm-wide acrylic cylinder with a spout for reward delivery (10% sucrose in water) in front of their mouth and a speaker located nearby. The behavioral apparatus was controlled via Arduino custom code. Reward delivery and licks measurement (capacitance sensor) was built on the model showed here (https://scanbox.org/2016/04/14/a-simple-lick-o-meter-and-liquid-reward-delivery-system/). Behavioral task events were transferred onto Excel with PLX-DAQ interface. Before the behavior started, mice were water-restricted to 85% of their body weight. Behavior started with 2 sessions of habituation, during which mice were headfixed and reward was randomly delivered through the spout. Then, conditioning started. Each session consisted of 50 trials. Each trial started with a 2s auditory cue (trilling 6kHz sound, 70dB), followed by a 1s delay period before reward delivery (Paired group). In the Unpaired group, the cue and reward were presented in a random order and never directly associated. Behavior happened in the dark, in a pseudo sound-isolated compartment, one session per day (total number of sessions specified in the figures). 15 minutes after the last training session, mice were anesthetized and slices were prepared for ex vivo recordings.

### Slice electrophysiology

The mice were anesthetized (ketamine/xylazine; 150 mg/100 mg kg^−1^), sacrificed, and their brains were transferred in ice-cold carbogenated (95% O_2_/5% CO_2_) solution, containing (in mM) choline chloride 110; glucose 25; NaHCO_3_ 25; MgCl_2_7; ascorbic acid 11.6; sodium pyruvate 3.1; KCl 2.5; NaH_2_PO_4_ 1.25; CaCl_2_0.5. Coronal brain slices (250 μm thickness) were prepared and transferred for 5 min to warmed solution (34°C) of identical composition, before transfer at room temperature in carbogenated artificial cerebrospinal fluid (ACSF) containing (in mM) NaCl 124; NaHCO_3_ 26.2; glucose 11; KCl 2.5; CaCl_2_ 2.5; MgCl_2_ 1.3; NaH_2_PO_4_ 1. During recordings, slices were immersed in ACSF and continuously superfused at a flow rate of 2.5 mL min^−1^ at 32°C. Neurons were patch-clamped using borosilicate glass pipettes (2.7–4 MΩ; Phymep, France) under an Olympus-BX51 microscope (Olympus, France). For voltage or current clamp recordings, signal was amplified, filtered at 5 kHz and digitized at 10 kHz (Multiclamp 200B; Molecular Devices, USA). Data were acquired using Igor Pro with NIDAQ tools (Wavemetrics, USA). Access resistance was continuously monitored with a −4 mV step delivered at 0.1 Hz. Experiments were discarded if the access resistance changed by more than 20% during the recording. Extracellular stimulation from AMPI ISO-Flex stimulator was delivered through glass electrodes placed in the LHb All recordings were made in voltage-clamp configuration, in APV-containing ACSF (100 μM). For E/I, evoked and spontaneous excitatory currents were recorded at −60mV, and evoked and spontaneous inhibitory currents were recorded at +5mV. Internal solution contained (in mM) cesium methanesulfonate 120, CsCl 10, HEPES 10, EGTA 10, creatine phosphate 5; Na2ATP 4; Na3GTP 0.4, QX-314 5. All drugs were purchased from HelloBio.

### Quantification and statistical analysis

Online and offline analyses of evoked synaptic currents were performed using Igor Pro-6 (Wavemetrics, USA). Spontaneous postsynaptic currents recordings were manually analyzed offline using Minianalysis (Synaptosoft Inc, USA). Sample size was predetermined on the basis of published studies and in-house expertise. Animals were randomly assigned to experimental groups. All data are represented as mean ±SEM and individual data points are shown. Data collection and analyses were not performed blinded to experimental conditions. Statistical comparisons were done in Prism (Graphpad) with the following methods: two-tailed Mann-Whitney test (for 2 independent groups); Kruskal-Wallis test followed by Dunn’s multiple comparison test (for three independent groups); 2-way ANOVA followed by Sidak’s test for multiple comparisons (Fig. 1d).

## Acknowledgements

We thank the entire Mameli Laboratory for discussions and comments on the manuscript and Ann Marie Craig for providing reagents. This work was supported by funds from the Canton of Vaud, the SNSF (31003A) and The Novartis Foundation to M.M.

## Author contributions

A.L.L. and M.M. conceptualized the project and wrote the manuscript with the help of all authors. A.L.L. performed and analyzed all experiments. J.A.C. and A.T. helped with *ex-vivo* recordings. M.C. helped with behavioral experiments. Y.G. designed and produced the Clptm1 tools for the project.

## Statement on data availability

The data sets generated during and/or analyzed during the current study are available from the corresponding author upon request.

---

**Supplementary Fig. 1.**
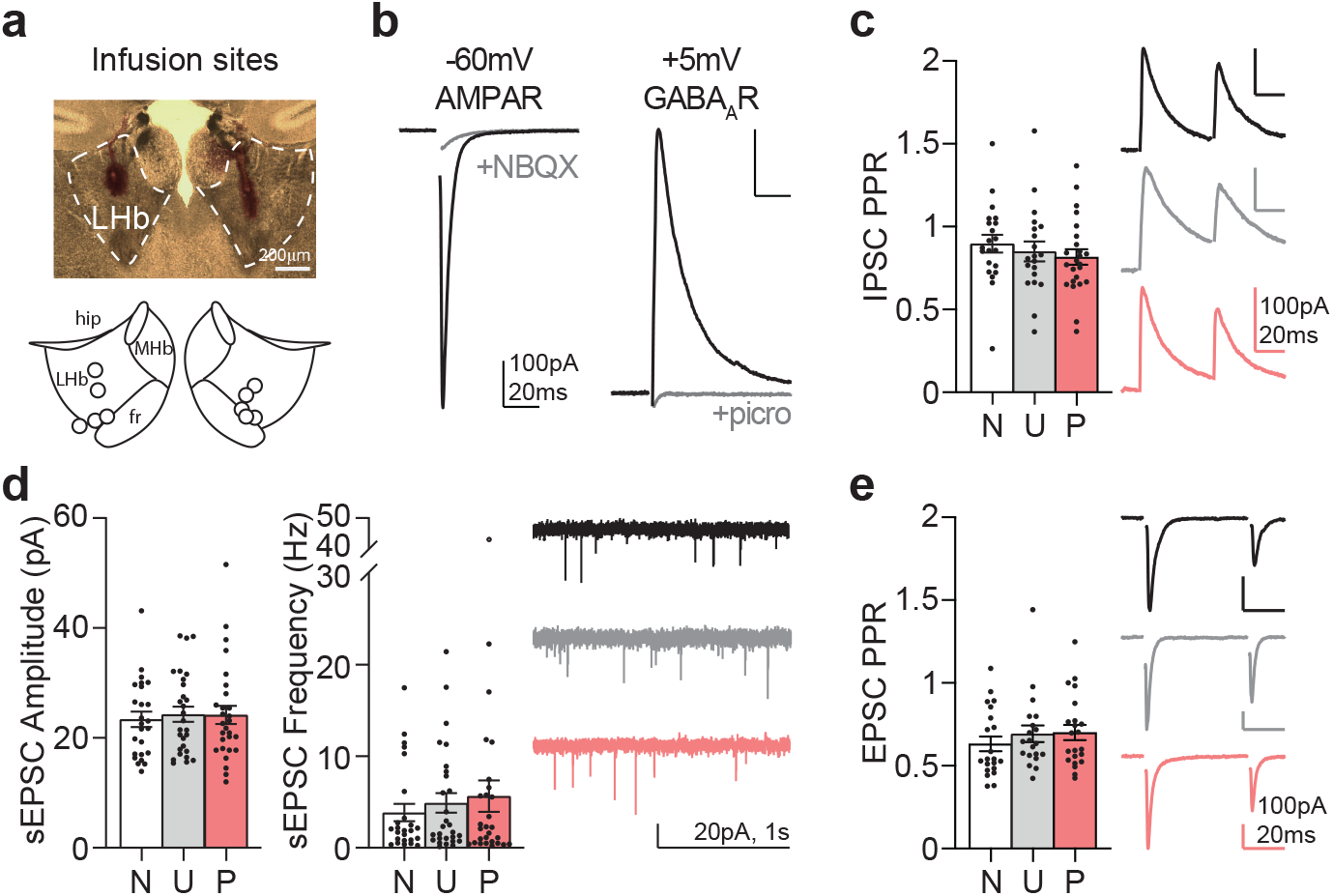
No change in presynaptic GABA release or excitatory transmission after conditioning. a. Example picture and infusion sites for picrotoxin/PBS. b. Example traces showing NBQX-sensitive inward current at −60mV and picrotoxin-sensitive outward current at +5mV. c. Paired-pulse ratio (PPR) of evoked inhibitory currents in naive (N, n=21), unpaired (U, n=20) and paired (P, n=24) mice (Kruskal-Wallis p=0.4675). c. Amplitude and frequency of excitatory spontaneous currents, and example traces (N, n=25, U, n=28, P, n=28; Kruskal-Wallis Amplitude p=0.8884; Frequency p=0.8582). d. PPR of evoked excitatory currents (N, n=21, U, n=20, P, n=22; Kruskal-Wallis p=0.2559)..

**Supplementary Fig. 2.**
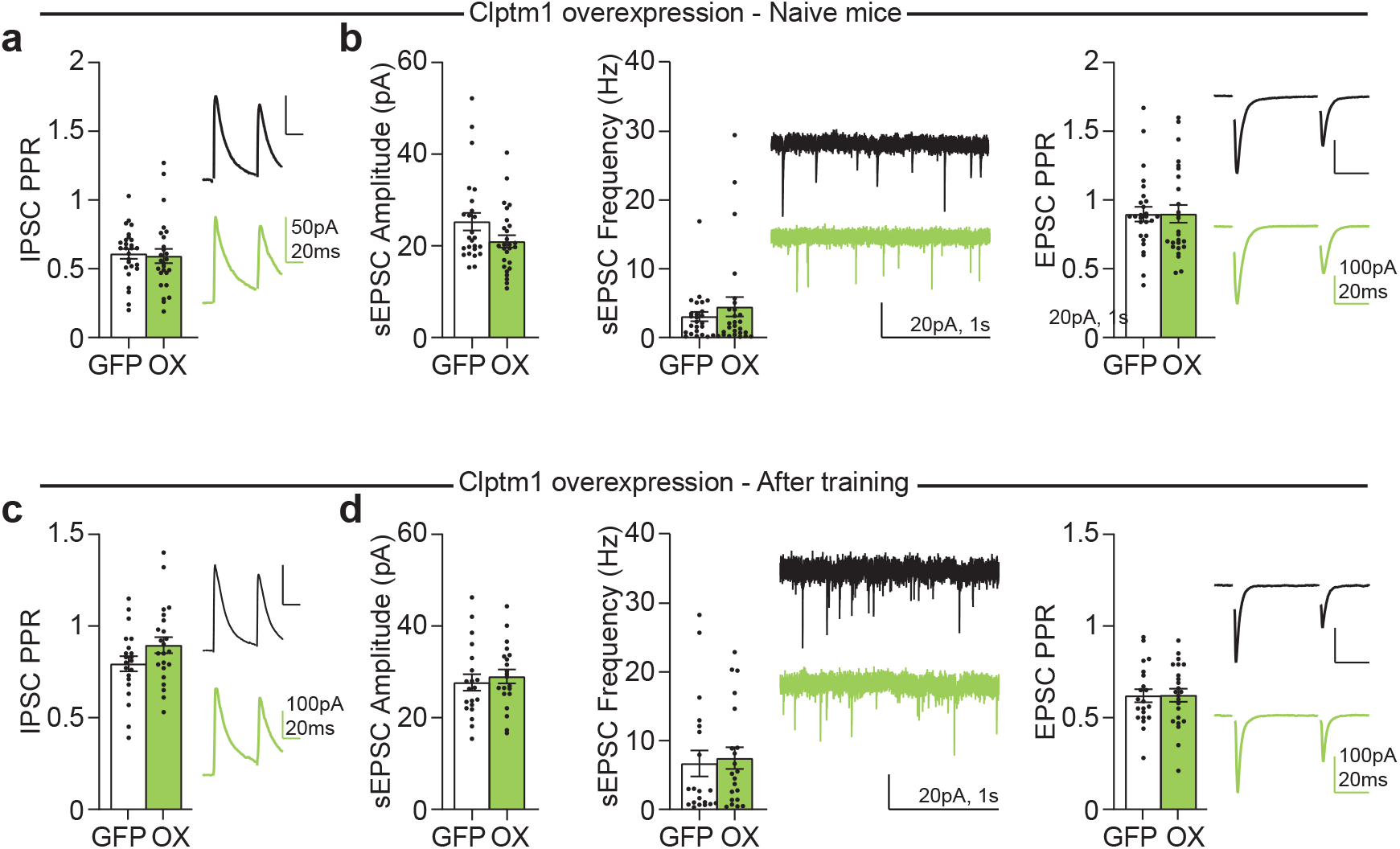
No change in presynaptic GABA release or excitatory transmission after Clptm1 overexpression. a. PPR of evoked inhibitory currents in naive mice overexpressing GFP (n=26) or Clptm1 (OX, n=26) in the LHb (p=0.4479). b. Amplitude and frequency of excitatory spontaneous currents (GFP, n=25, OX, n=27, Amplitude p=0.1202, Frequency p=0.7541) and PPR of evoked excitatory currents in naive mice (GFP, n=26, OX, n=26, p=0.1907). c. PPR of evoked inhibitory currents after Pavlovian conditioning (GFP, n=21, OX, n=23, p=0.8935). d. Amplitude and frequency of excitatory spontaneous currents (GFP, n=20, OX. N=21, Amplitude p=0.4007, Frequency p=0.4935) and PPR of evoked excitatory currents after Pavlovian conditioning (GFP, n=21, OX, n=23, p=0.1580).

